# VSIG4-Expressing Macrophages Contribute To Anti-Parasitic And Anti-Metastatic Responses In The Peritoneal Cavity

**DOI:** 10.1101/2024.12.23.630094

**Authors:** Els Lebegge, Daliya Kancheva, Jolien Van Craenenbroeck, Sam Ernst, Pauline M.R. Bardet, Aarushi A. Caro, Maté Kiss, Neema Ahishakiye Jumapili, Romina Mora Barthelmess, Maida Zivalj, Naela Assaf, Yvon Elkrim, Jesse Demuytere, Jan De Jonge, Geert Raes, Eva Hadadi, Nick Devoogdt, Cécile Vincke, Lars Vereecke, Wim Ceelen, Benoit Stijlemans, Damya Laoui, Sana M. Arnouk, Jo A. Van Ginderachter

## Abstract

Peritoneal tissue-resident macrophages, also referred to as large peritoneal macrophages (LPMs), play an important role as gatekeepers of peritoneal homeostasis by providing a first line of defense against pathogenic threats. About a third of the LPMs express the surface receptor V-set and Immunoglobulin domain containing 4 (VSIG4), but it is unclear to what extent these cells differ from their VSIG4-negative counterparts and perform dedicated functions. Here, we demonstrate that VSIG4^+^ LPMs, in contrast to VSIG4^-^ LPMs, are in majority derived from embryonal precursors and their occurrence is to a large extent independent from sex and microbiota. Although their transcriptome and surface proteome are indistinguishable from VSIG4^-^ LPMs at steady-state, VSIG4^+^ LPMs are superior in phagocytosing Gram-positive bacteria and colorectal carcinoma (CRC) cells. In-house generated anti-VSIG4 nanobody constructs that are antibody-dependent cell-mediated cytotoxicity (ADCC)-enabled allowed a selective elimination of the VSIG4^+^ LPM subset without affecting the overall LPM content of the peritoneal cavity. This strategy uncovered a role for VSIG4^+^ LPMs in lowering the first peak of parasitemia in a *Trypanosoma brucei brucei* infection model and in reducing the outgrowth of CRC cells in the peritoneal cavity, a prime metastatic site in CRC patients. Altogether, our data uncover a protective role for VSIG4^+^ LPMs in infectious and oncological diseases in the peritoneal cavity.

## INTRODUCTION

In the peritoneal cavity (PC) at steady-state, two subsets of tissue-resident macrophages have been described based on their morphology and function, termed “small” and “large” peritoneal macrophages (SPMs and LPMs)^1^. Disturbance of homeostasis in the PC leads to LPM activation, followed by their adhesion and cellular clot formation, disappearance from the PC and migration to the omentum^2^. Of interest, a subset of LPMs expresses the V-set and Immunoglobulin domain containing 4 receptor (VSIG4), which is a member of the B7-protein family and is also referred to as complement receptor of the immunoglobulin superfamily (CRIg)^3,4^. However, it is currently unclear to what extent the VSIG4^+^ and VSIG4^-^ LPM subsets differ and whether they have dedicated functions.

VSIG4 ligand binding, which leads to receptor internalization, can occur in a complement-dependent, e.g. C3-opsonized particles, and in a complement-independent manner, e.g. lipoteichoic acid binding on Gram-positive bacteria^3,5^. VSIG4 receptor signaling in macrophages suppresses their pro-inflammatory activation through metabolic changes, leading to reduced mitochondrial reactive oxygen species production^6^. Besides these macrophage-intrinsic effects, the VSIG4 receptor can also suppress T-cell activation and T-cell-mediated responses, either by directly binding to an as yet unidentified receptor on T cells or via complement proteins that bridge the VSIG4 receptor and a complement receptor on T cells^4,7,8^. Hence, the VSIG4 receptor is able to modify both innate (macrophage-mediated) and adaptive immunity and is therefore an interesting target for the treatment of macrophage-driven pathologies^9^.

For this reason, we explored the potential differences between VSIG4^+^ and VSIG4^-^ LPMs, the function of the VSIG4 receptor, as well as the possibility to target VSIG4^+^ LPMs. scRNA-seq and CITE-seq data showed a lack of transcriptomic and proteomic differences between VSIG4^+^ and VSIG4^-^ LPMs at steady-state and during an infection with African trypanosomes, but VSIG4^+^ LPMs demonstrated an increased phagocytic capacity. A specific depletion of VSIG4^+^ LPMs via a newly generated anti-VSIG4-antibody-dependent cell-mediated cytotoxicity (ADCC) construct led to an enhanced first peak of *Trypanosoma* parasitemia and a faster peritoneal dissemination of colorectal carcinoma (CRC) cancer cells, showing a protective function of this peritoneal macrophage population in such disease settings.

## RESULTS

### VSIG4 marks a subset of large peritoneal macrophages (LPMs) that is ontogenically different from VSIG4-negative LPMs and is maintained regardless of sex and microbiota

To assess VSIG4 expression on the PC CD45^+^ hematopoietic compartment, a lavage from the steady-state PC of female C57BL/6 mice was analyzed. LPMs can be identified through their high F4/80 and ICAM2 expression (Fig 1A), with approximately one third of these cells expressing high levels of the VSIG4 receptor. Conversely, VSIG4 was absent on SPMs and other CD45^+^ cell types in a naive peritoneal lavage (Fig 1B). In age-matched male mice, the overall percentage of LPMs in the PC was not significantly different from females (Fig S1A), but the percentage of VSIG4^+^ cells within that population was on average 5% lower (Fig S1B). Of note, the VSIG4 surface expression level was not significantly different between male and female VSIG4^+^ LPMs, but tended to be lower in the latter (Fig S1C).

**Figure 1.**
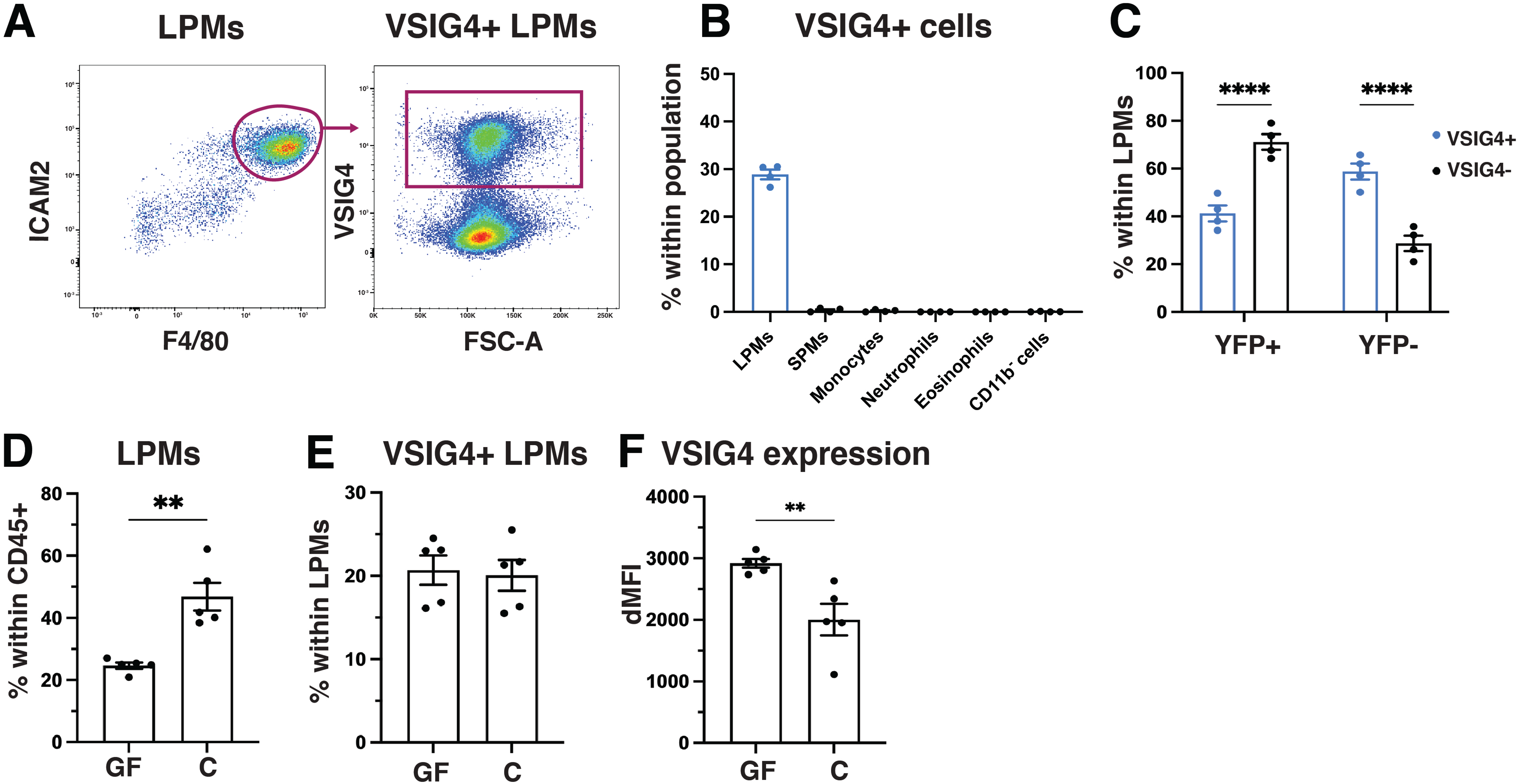
VSIG4 marks a subset of large peritoneal macrophages (LPMs) in the peritoneal cavity. **(A)** Representative gating strategy for the identification of VSIG4^+^ LPMs in a naive peritoneal lavage of female C57BL/6 mice. **(B)** Percentage of VSIG4^+^ cells within each immune cell population in a naive peritoneal lavage of female C57BL/6 mice (n=4). **(C)** Percentage of YFP^+^ and YFP^-^ cells within VSIG4^+^ and VSIG4^-^ LPMs of naive Flt3-Cre x ROSA26-YFP mice (n=4). **(D)** Percentage of LPMs within CD45^+^ cells, **(E)** percentage of VSIG4^+^ LPMs within all LPMs and **(F)** VSIG4 expression level in VSIG4^+^ LPMs depicted as ΔMFI in naive germ-free C57BL/6 mice compared to conventional mice (n=5/group). Data is shown as the mean ± SEM.

Like many tissue-resident macrophages, the LPM pool is sustained via the proliferation of embryonic LPMs, but can also be replenished by bone marrow-derived monocytes^10,11^. To assess the ontogeny of VSIG4^+^ and VSIG4^-^ LPMs, Flt3-driven YFP-reporter mice were used, in which YFP^+^ cells are of bone marrow monocytic origin^12^. Interestingly, approximately 60% of VSIG4^+^ LPMs were YFP-negative and hence of embryonic origin, while the majority (70%) of VSIG4^-^ LPMs were YFP-positive and hence of bone marrow origin (Fig 1C). Thus, although VSIG4 expression is mostly associated with embryonic LPMs, this receptor can also be acquired by monocyte-derived cells.

Since VSIG4 functions as a pattern-recognition receptor, the presence of bacterial molecules could be another factor regulating VSIG4 expression by LPMs. Indeed, previous work demonstrated an impact of the microbiota on peritoneal macrophages^13^, prompting us to ask whether the gut microbiome could affect VSIG4^+^ LPMs. In germ-free (GF) mice, the percentage of LPMs within CD45^+^ peritoneal cells was significantly lower as compared to age-matched conventional mice (Fig 1D), but within the LPM population an equal contribution of VSIG4^+^ cells was observed (Fig 1E). However, the VSIG4 expression level was significantly higher on VSIG4^+^ LPMs of GF mice (Fig1F), suggesting that intestinal microbes or their derived factors may restrain VSIG4 surface expression but do not fundamentally affect the generation of VSIG4^+^ LPMs.

### Steady-state VSIG4^+^ and VSIG4^-^ LPMs are identical at the transcriptomic and surface proteomic level, but VSIG4^+^ LPMs are superior phagocytes of Gram-positive bacteria

Since VSIG4 signaling is able to affect the macrophage activation state^6^, it is conceivable that VSIG4-expressing LPMs may be differently activated from their VSIG4-negative counterparts. To evaluate differences between steady-state VSIG4^+^ and VSIG4^-^ LPMs at the transcriptomic level, LPMs were in first instance extracted from the publicly available dataset by Bain et al^12^, encompassing peritoneal cells from female and male mice, based on the expression of tissue-resident LPM markers (*C1qa, Cd9, Gata6, Icam2, Timd4, Prg4, Calml4, Prtn3*) and exclusion of clusters with T cell (*Cd3e, Cd3g, Cd2*), monocyte (*Ccr2, Ly6c2, Plac8, Ear2*) and DC (*Flt3, H2-Aa, Itgax, Cd209a, Slamf7, Pldb1*) specific genes. Four LPM clusters were obtained after excluding proliferating cells from the scRNAseq dataset (Fig 2A), with all clusters expressing variable levels of prototypical LPM genes such as *Adgre1, Timd4, Icam2, Cd9, C1qc, C1qb* and *Cd68* (Fig 2B). These LPM genes were most prominently expressed by the small cluster 3, that also expresses the highest levels of *Marco* (Fig 2B). Cluster 1 most prominently expressed *Sdc3, Wfdc17, Atp5e* and *Rpl22l1* and cluster 2 was characterized by *Folr2, Ccl6, Retnla, Mrc1, Ccr2* and genes linked to MHC-II expression (*Cd74, H2-Aa, H2-Ab1*) (Fig 2B). Remarkably, *Vsig4* expression was spread across all LPM subsets, all of which were partially *Vsig4*^+^, with the lowest expression level seen in cluster 2 and the highest in cluster 0 (Fig 2B,C). When the entire LPM population was separated into a *Vsig4*^+^ and a *Vsig4*^-^ subset, no significantly differentially expressed genes (DEGs, with a fold change higher than 1) were discovered, except for the *Vsig4* gene itself (Fig 2D). Of note, when analyzing samples from female and male mice separately, the same LPM subsets were detected in both sexes (Fig S2A-B), no transcriptomic differences were found between female and male *Vsig4^+^* LPMs (Fig S2C) and only a few genes were differentially expressed between *Vsig4^-^* LPMs in both sexes (Fig S2D). Moreover, no DEGs (except *Vsig4*) were found between *Vsig4^+^* and *Vsig4^-^* LPMs, either in females or males (Fig S2E-F).

**Figure 2.**
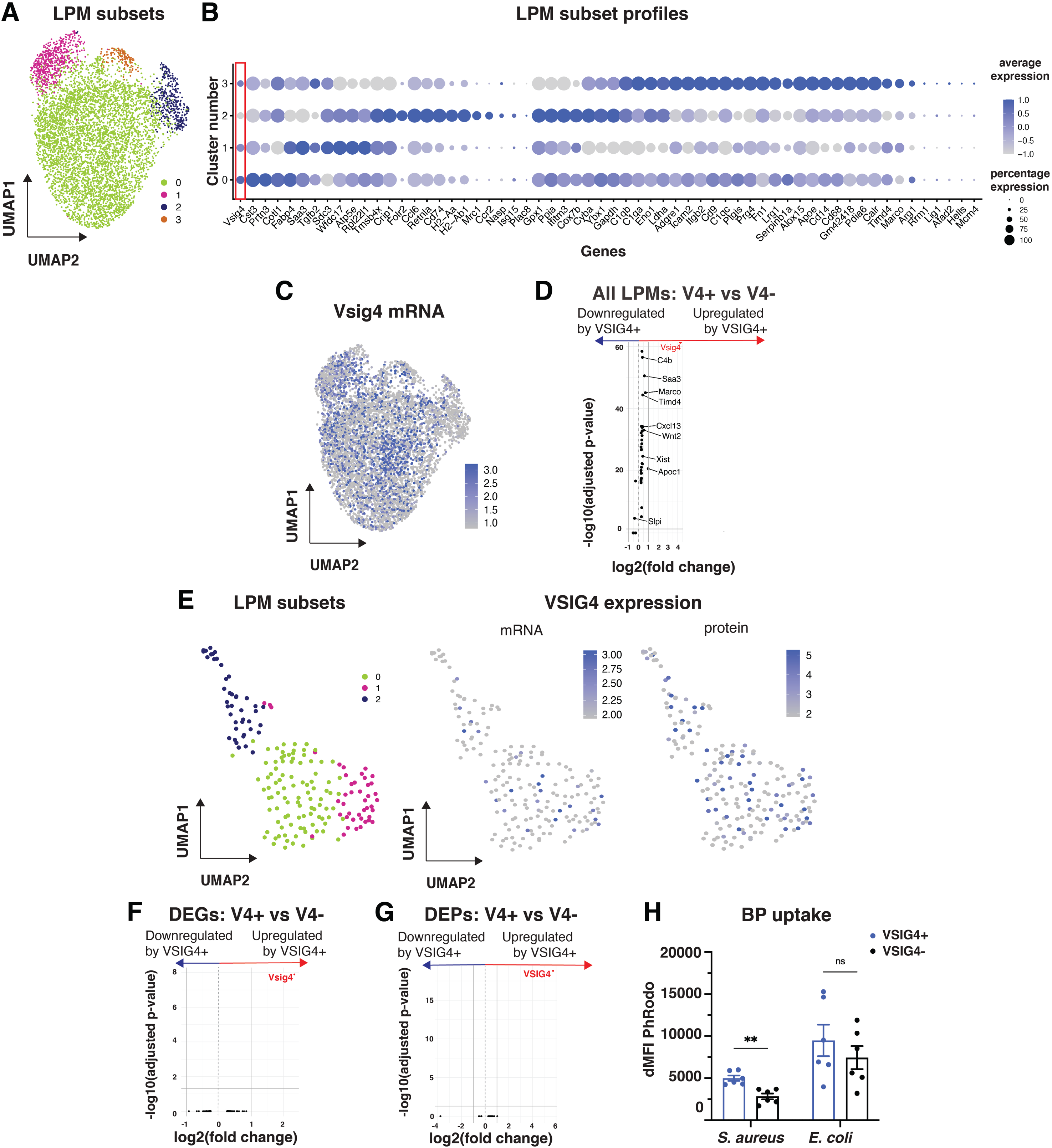
VSIG4^+^ LPMs are superior phagocytes for Gram-positive bacteria, despite being identical at the transcriptomic and surface proteomic level in steady state. **(A)** UMAP plot of LPM clusters obtained from the scRNA-sequencing dataset by Bain et al^26^. **(B)** Dot plot of highly expressed markers per LPM subset. The size of the dot indicates the expression percentage in the LPM subset, while the color indicates the average expression level. **(C)** Feature plot of *Vsig4* mRNA expression in LPM subsets. **(D)** Volcano plot of differentially expressed genes (DEGs) between *Vsig4*^+^ and *Vsig4*^-^ LPMs, whereby the adjusted p-value is plotted versus the fold change. LPMs were divided into *Vsig4*^+^ and *Vsig4*^-^ based on the mRNA expression cut-off of >2 and 0, respectively. **(E)** UMAP plot of LPM clusters, obtained via CITE-sequencing of peritoneal exudate cells obtained from control i.p. HBSS-injected mice (left), and feature plot of *Vsig4* mRNA and VSIG4 protein expression in LPM subsets (right). Volcano plot of DEGs **(F)** and differentially expressed proteins (DEPs) **(G)** between VSIG4^+^ and VSIG4^-^ LPMs, whereby the adjusted p-value is plotted versus the fold change. LPMs were divided into VSIG4^+^ and VSIG4^-^ LPMs based on the protein expression cut-off of >3.5 and 0, respectively. **(H)** *In vivo* phagocytosis of PhRodo-labeled *S. aureus* and *E. coli* bioparticles, depicted as ΔMFI of the PhRodo signal in VSIG4^+^ and VSIG4^-^ LPMs. Unlabeled *S. aureus* bioparticles and HBSS were used as negative controls, respectively (n=5/group). Data is shown as the mean ± SEM.

To assess potential differences between steady-state VSIG4^+^ and VSIG4^-^ LPMs at the surface proteome level, a cellular indexing of transcriptomes and epitopes (CITE)-sequencing analysis on peritoneal lavages of female C57BL/6 mice was performed (Table S2 for the list of barcoded antibodies), incorporating barcoded anti-VSIG4 antibodies^14^. This newly generated dataset again revealed LPM heterogeneity based on transcriptomic differences (Fig 2E, Fig S3) and, similar to the *Vsig4* mRNA expression pattern (Fig S3), the VSIG4 protein was expressed at the surface across all different LPM subsets (Fig 2E). Consequently, when LPMs were split into VSIG4^+^ and VSIG4^-^ subsets, hardly any differentially expressed genes (DEGs) nor proteins (DEPs) were detected between naive VSIG4^+^ and VSIG4^-^ LPMs (Fig 2F,G). In conclusion, at steady-state, no transcriptomic nor surface proteomic differences can be observed between VSIG4^+^ and VSIG4^-^ LPM subsets, except VSIG4 itself.

Finally, we assessed whether the expression of VSIG4, which can serve as a pattern recognition receptor by binding lipoteichoic acid in the cell wall of Gram-positive bacteria such as *Staphylococcus aureus*^5^, confers a superior bacterial uptake capacity to LPMs. To this end, the phagocytic capacity of VSIG4^+^ and VSIG4^-^ LPMs towards PhRodo-labeled *S. aureus* and Gram-negative *E. coli* bioparticles (BP) was tested, by injecting these BP in the PC of naïve female C57BL6 mice and collecting peritoneal lavages 10 minutes later. A significantly increased uptake of *S. aureus* BP, but not *E. coli* BP, was observed by VSIG4^+^ LPMs as compared to VSIG4^-^ LPMs (Fig 2H). Hence, despite being identical at steady-state, VSIG4 expression confers a competitive advantage to VSIG4^+^ LPMs to phagocytose Gram-positive bacteria.

### A specific nanobody-mediated depletion of VSIG4^+^ LPMs demonstrates their influence on early phase *Trypanosoma brucei brucei* infections without effect on chronic disease outcome

VSIG4 is also known to play a role in the phagocytic uptake of African trypanosomes, such as *Trypanosoma brucei brucei*^19^. Therefore, we injected live *T. b. brucei* parasites in the PC and assessed the response of LPMs 18h later. At that timepoint, the percentage of LPMs within the total CD45^+^ population tended to be reduced in infected as compared to naïve mice (Fig 3A), likely due to the well-known “macrophage disappearance reaction”^2,15^. Nevertheless, the relative contribution of VSIG4^+^ cells to the overall LPM pool remained unaltered (Fig 3B), although the VSIG4 surface expression level on VSIG4^+^ LPMs went down in infected mice (Fig 3C), possibly reflecting receptor internalization upon binding. CITE-sequencing uncovered novel LPM clusters in infected mice as compared to the naïve controls. Indeed, while cells belonging to clusters 0, 1 and 2 could be found in both naive and infected mice, clusters 3 (characterized by *Cox7b*, *Ifitm3*, *Pycard* and *Tmsb4x*) and 4 (characterized by *Ccl6, Saa3, Sdc3, Ldha* and *Arg1*) were almost exclusively found in the infected PC (Fig 3D,E; Fig S4). Upon reclustering of LPMs from *T. brucei brucei*-infected mice only, *Vsig4* mRNA as well as VSIG4 protein were expressed across all LPM subsets, akin to the naïve situation (Fig 3F). An increased gene expression of *Ccl6*, *Saa3*, *Arg1* and *Zbp1* was detected in VSIG4^+^ LPMs of infected mice compared to naïve VSIG4^+^ LPMs (Fig 3G), suggestive of a somewhat more inflammatory phenotype^16–19^. In accordance, at the protein level, VSIG4^+^ LPMs of infected mice showed an upregulation of proteins related to macrophage activation (MHC-II, CD172a, CD55, Ly6A), migration (CD105, CD102, CD63), as well as modulation of the adaptive immune response (CD80, CD86, CD274, CD150) as compared to naïve VSIG4^+^ LPMs (Fig 3H). However, no significant DEGs nor DEPs, except for *Vsig4/*VSIG4, were detected between VSIG4^+^ and VSIG4^-^ LPMs in the infected PC (Fig 3I,J).

**Figure 3.**
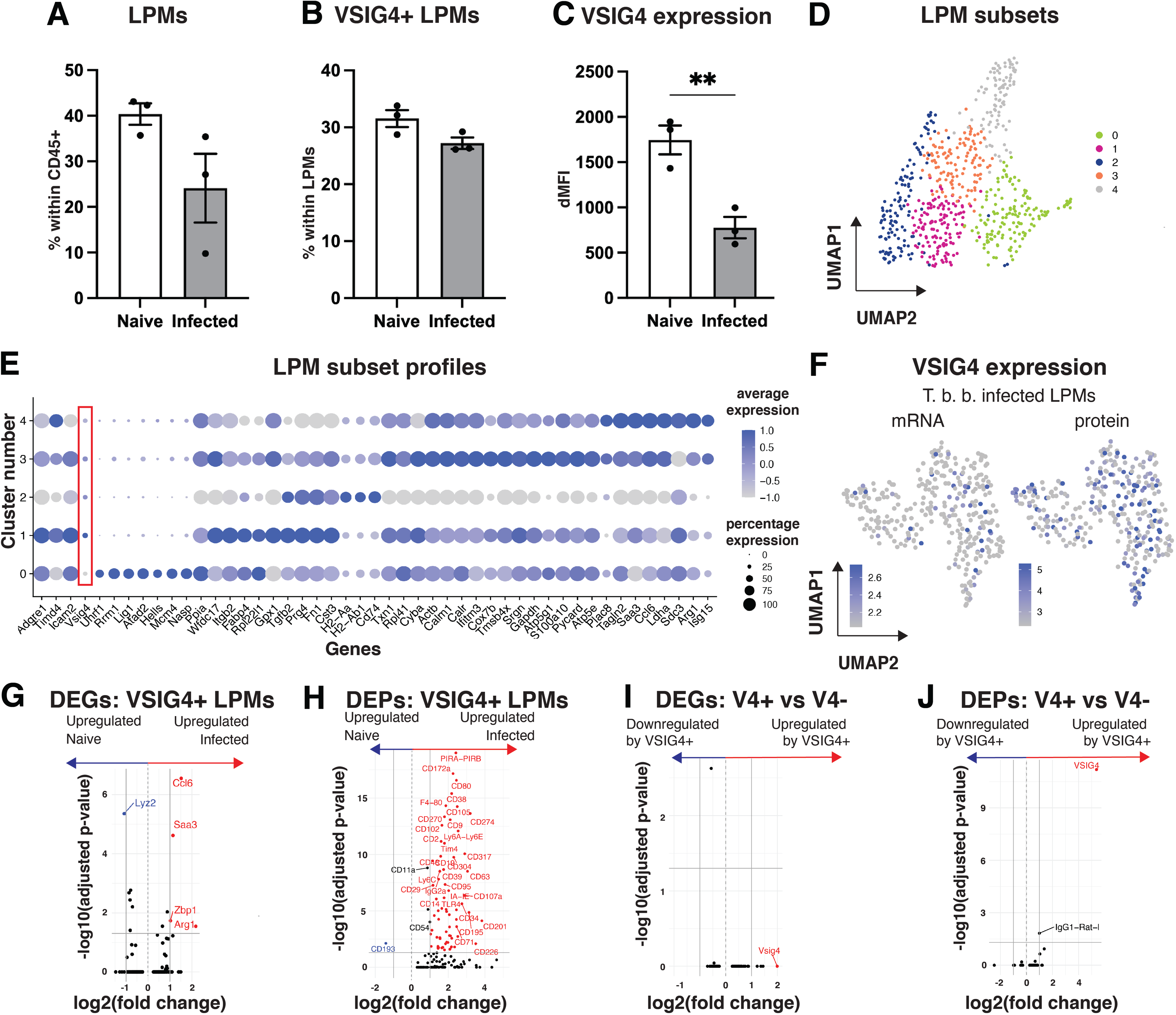
VSIG4^+^ LPMs influence the early phase of *Trypanosoma brucei brucei* infections. **(A)** Percentage of LPMs within CD45^+^ cells, **(B)** Percentage of VSIG4^+^ LPMs within all LPMs and **(C)** VSIG4 expression level in VSIG4^+^ LPMs depicted as ΔMFI in peritoneal lavages of female C57BL/6 mice 18 hours post i.p. injection of *Trypanosoma brucei brucei* or HBSS (naive) (n=3/group). **(D)** UMAP plot of LPM clusters obtained via CITE-sequencing of peritoneal exudate cells from HBSS-injected and *T. brucei brucei*-infected mice. **(E)** Dot plot of highly expressed markers per LPM subset in the combined HBSS-injected and *T. brucei brucei*-infected dataset. The size of the dot indicates the expression percentage in the LPM subset, while the color indicates the average expression level. **(F)** Feature plot of *Vsig4* mRNA and VSIG4 protein expression in LPM subsets of *T. brucei brucei*-infected mice. Volcano plot of DEGs **(G)** and DEPs **(H)** between VSIG4^+^ LPMs of HBSS-injected and *T. brucei brucei-*infected mice, and **(I)** DEGs and **(J)** DEPs between VSIG4^+^ and VSIG4^-^ LPMs of *T. brucei brucei-*infected mice, whereby the adjusted p-value is plotted versus the fold change. LPMs were divided into VSIG4^+^ and VSIG4^-^ LPMs based on the protein expression cut-off of >3.5 and 0, respectively. Data is shown as the mean ± SEM.

To directly investigate the role of VSIG4^+^ LPMs during a *T. b. brucei* infection, we developed a strategy for the specific elimination of this LPM subset. An anti-VSIG4-ADCC construct was designed by fusing an ADCC-enabled mouse IgG2a Fc tail to two copies of a mouse VSIG4-binding VHH, that was previously generated by our lab^17^ (Fig 4A). As a non-depleting control, an ADCC-disabled construct was generated by introducing a LALAPG-mutation in the IgG2a Fc-tail^21^. Seven days after a single intraperitoneal injection of the anti-VSIG4-ADCC construct in naïve female animals, the majority of VSIG4^+^ LPMs was depleted, while this was not the case with the ADCC-disabled construct (Fig 4B). VSIG4^+^ LPMs remained largely absent for at least another two weeks. Importantly, the percentage of total LPMs in the PC remained unaltered upon depletion, suggesting a rapid replenishment of the total LPM pool, albeit without VSIG4 expression (Fig 4C). As a matter of fact, VSIG4^+^ LPMs did not fully recover by day 56 post-depletion, illustrating that regaining the VSIG4^+^ phenotype is a slow process (Fig 4B). Hence, a single intraperitoneal injection of anti-VSIG4-ADCC allows the generation of a peritoneal cavity that lacks VSIG4^+^ LPMs, without affecting the total percentage of LPMs.

**Figure 4.**
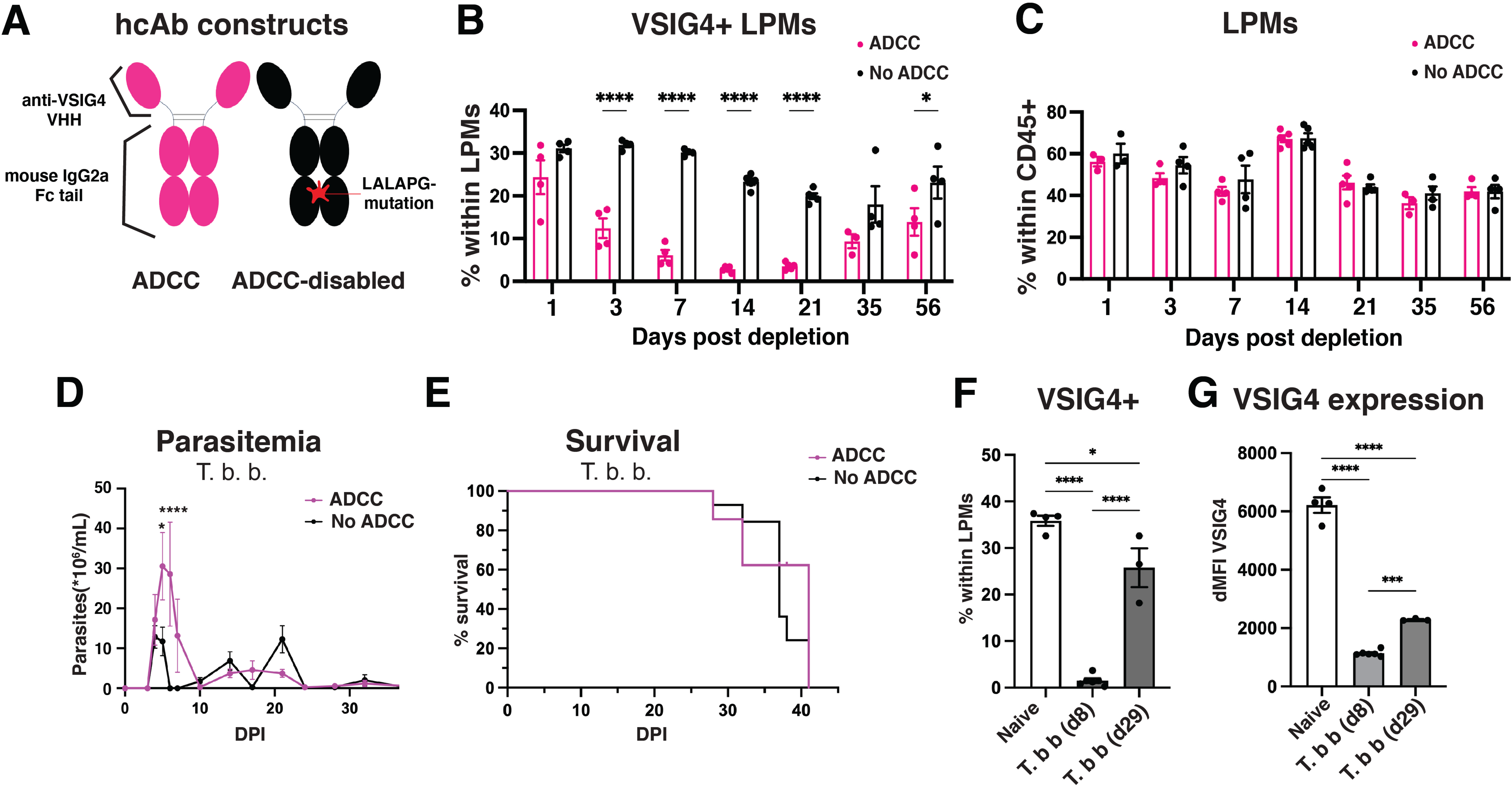
Depletion of VSIG4^+^ LPMs during a *Trypanosoma brucei brucei* infection increases the first peak of parasitemia. **(A)** Schematic representation of the ADCC-enabled (pink) and ADCC-disabled (black) anti-VSIG4 constructs. **(B)** Percentage of VSIG4^+^ LPMs within all LPMs and **(C)** percentage of LPMs within CD45^+^ peritoneal exudate cells of naive C57BL/6 mice at different intervals following a single i.p. injection of the anti-VSIG4 ADCC-enabled (pink) or ADCC-disabled (black) construct (n=4/group). **(D)** Parasitemia in the blood and **(E)** Kaplan-Meier survival curve of C57BL/6 mice that received a single i.p. injection of the anti-VSIG4 ADCC-enabled (pink) or ADCC-disabled (black) construct 7 days prior to the *T. brucei brucei*-infection (n=7/group). **(F)** Percentage of VSIG4^+^ LPMs within all LPMs and **(G)** VSIG4 expression level in VSIG4^+^ LPMs depicted as ΔMFI, in naive (n=4), 8 days post-*T. brucei brucei* infection (n=6) and 29 days post-infection (n=3) C57BL/6 mice. Data is shown as the mean ± SEM.

Using this strategy, VSIG4^+^ LPMs were depleted seven days prior to the *T. b. brucei* infection. At d5 and d6 post-infection, an increased peak parasitemia was observed in mice that received the VSIG4-depleting construct (Fig 4D), suggesting the contribution of VSIG4^+^ LPMs to early parasite elimination. However, no increased parasitemia was observed at later stages and the mice survived equally long (Fig 4E). The lack of long-term effects of experimental VSIG4^+^ LPM depletion on infection parameters is in line with the fact that *T. b. brucei* naturally downscales the presence of these cells. Indeed, in female C57BL/6 hosts, the VSIG4^+^ LPM population nearly disappeared at d8, to return to approximately 25% of the total LPM population by d29 (Fig 4F), albeit with a strongly reduced VSIG4 surface expression level (Fig 4G). Hence, VSIG4^+^ LPMs contribute to parasite elimination in the early phase of infection.

### VSIG4^+^ LPMs are superior in phagocytosing CRC cancer cells and protect against CRC peritoneal tumor outgrowth

Since the PC is a common metastatic site for colorectal carcinoma (CRC)^22,23^, it can be questioned whether VSIG4^+^ LPMs play a role in the establishment and growth of CRC peritoneal metastases.

Human peritoneal macrophages were shown before to express VSIG4 in healthy conditions^24^. We now assessed whether the VSIG4 receptor is expressed on macrophages associated with peritoneal metastatic nodules of CRC cancer patients, in comparison to their primary tumor. Macrophages were identified as CD45^+^ CD11b^+^ CD66b^-^ CD14^+^ HLA-DR^+^ CD163^+^ cells (Fig S5) and their percentage (within CD45^+^) was on average higher in CRC metastatic nodules in the omentum (O) and mesentery (M) as compared to the primary tumor (PT) (Fig 5A). Importantly, VSIG4^+^ macrophages were present in both the metastatic nodules and the primary tumor (Fig 5B), suggesting a potential involvement of these cells in the metastatic process.

**Figure 5.**
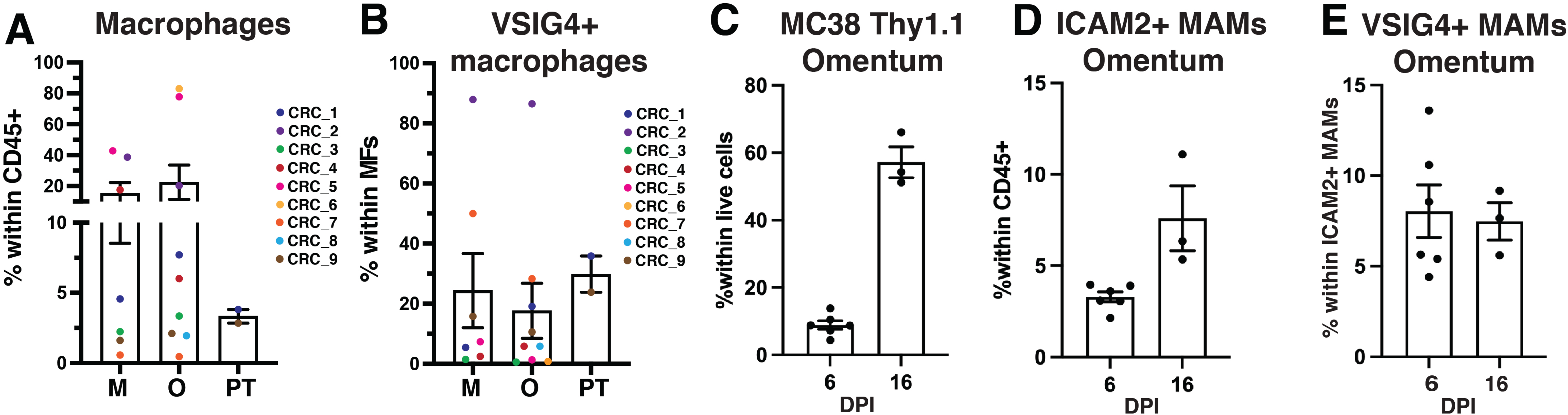
VSIG4^+^ macrophages are present in peritoneal metastasis of CRC patients and a murine model of CRC metastasis. **(A)** Percentage of macrophages within CD45^+^ cells and **(B)** VSIG4^+^ macrophages within all macrophages in the primary tumor (PT) and metastasis to the mesentery (M) and omentum (O) of 9 patients with metastasized colorectal cancer. **(C)** Percentage of MC38-Thy1.1 cells within all live cells in the omentum, **(D)** percentage of ICAM2^+^ metastasis-associated macrophages (MAMs) within CD45^+^ cells and **(E)** percentage of VSIG4^+^ MAMs within all MAMs following 6 days (n=6) and 16 days (n=3) post-MC38-Thy1.1 injection (days post injection, DPI) in C57BL/6 mice. Data is shown as the mean ± SEM.

To investigate the role of VSIG4^+^ LPMs in a mouse model of CRC peritoneal metastasis, mouse MC38-Thy1.1^+^ CRC cancer cells were injected in the PC. In this model, MC38-Thy1.1^+^ nodules were present in the omentum as early as 6 days post-injection, which further increased by day 16 (Fig 5C). These nodules contained ICAM2^+^ metastasis-associated macrophages (MAMs), a percentage of which also expressed the VSIG4 receptor (Fig 5D,E). In a first setting, VSIG4^+^ LPMs were depleted by injecting the anti-VSIG4-ADCC construct 7 days prior to MC38-Thy1.1^+^ cancer cell injection (Fig 6A), at which point the cancer cells are confronted with a PC largely devoid of VSIG4^+^ LPMs while the total LPM pool was unaltered (Fig 4B,C). As a control, the anti-VSIG4-LALAPG ADCC-disabled construct was injected. Prior depletion of VSIG4^+^ LPMs resulted in a significantly decreased survival as compared to control mice, suggesting that VSIG4^+^ LPMs play a protective role during MC38 growth in the PC (Fig 6B). To assess the effect of VSIG4^+^ LPM depletion when tumors were initiated, we treated mice with anti-VSIG4-ADCC or the non-depleting control construct two days after the intraperitoneal cancer cell inoculation (Fig 6C). Again, the depletion of VSIG4^+^ LPMs resulted in a decreased survival as compared to control mice (Fig 6D). Together, these data demonstrate a protective role of VSIG4^+^ LPMs against CRC tumor outgrowth in the PC.

**Figure 6.**
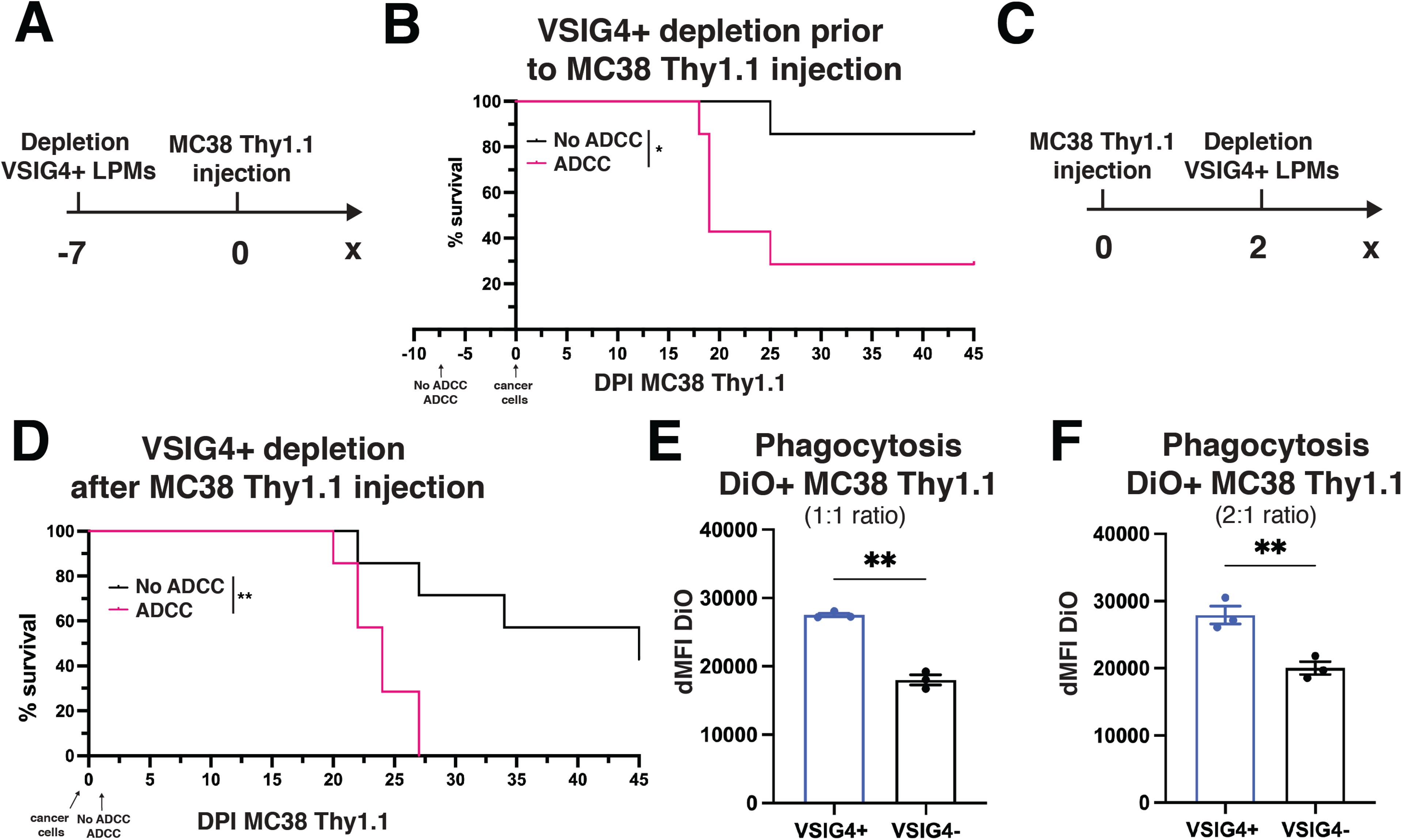
VSIG4^+^ LPMs are superior in phagocytosing CRC cancer cells and protect against CRC peritoneal metastasis. **(A)** Schematic representation of the cancer cell and anti-VSIG4 construct injection regimen of the experiment shown in panel B. **(B)** Kaplan-Meier survival curve of C57BL/6 mice that received an i.p. injection of the anti-VSIG4 ADCC-enabled (pink) or ADCC-disabled (black) construct 7 days prior to MC38-Thy1.1 cancer cell injection (n=7/group). **(C)** Schematic representation of the cancer cell and anti-VSIG4 construct injection regimen of the experiment shown in panel D. **(D)** Kaplan-Meier survival curve of C57BL/6 mice that received an intraperitoneal injection of the anti-VSIG4 ADCC-enabled (pink) or ADCC-disabled (black) construct 2 days after MC38-Thy1.1 cancer cell injection (n=7/group). *In vitro* phagocytosis of DiO-labeled MC38-Thy1.1 cancer cells, **(E)** at a ratio of 1:1 (peritoneal exudate cells:cancer cells), or **(F)** at a ratio of 2:1 (peritoneal exudate cells : cancer cells), depicted as ΔMFI of the DiO signal in VSIG4^+^ and VSIG4^-^ LPMs. Unlabeled MC38-Thy1.1 cancer cells were used as negative control for the DiO signal (n=3/group). Data is shown as the mean ± SEM.

To mechanistically underpin these findings, we assessed the phagocytic capacity of VSIG4^+^ and VSIG4^-^ LPMs towards MC38-Thy1.1^+^ cancer cells. Hereto, DiO-labeled MC38-Thy1.1^+^ cancer cells were co-cultured for 1h with peritoneal exudate cells, after which the DiO-positivity of VSIG4^+^ and VSIG4^-^ LPMs was assessed via flow cytometry. Interestingly, VSIG4^+^ LPMs displayed a significantly enhanced capacity to phagocytose MC38-Thy1.1^+^ cancer cells as compared to VSIG4^-^ LPMs (Fig 6E,F), possibly explaining their importance as anti-tumoral cells in the PC.

## DISCUSSION

To date, the function of the VSIG4 receptor has been mostly described in the liver, where it is expressed by the entire liver-resident Kupffer cell (KC) population and is involved in complement-dependent and -independent pathogen recognition. We and others showed that approximately 30% of the LPMs in the PC also expresses the VSIG4 receptor, but its regulation and function on those cells, as well as the significance of VSIG4^+^ LPMs during steady-state or peritoneal infectious or cancerous disease is unknown.

A remarkable finding is that the majority of VSIG4^+^ LPMs was of embryonic origin, while most of the VSIG4^-^ LPMs were of monocytic origin. This could possibly be explained by a steady turnover of embryonically derived LPMs by monocyte-derived LPMs, whereby VSIG4 expression is only slowly gained by the monocyte-derived cells. In accordance, Louwe et al demonstrated that the time-of-residency in the PC is a determining factor for the expression of LPM markers such as VSIG4, Tim4 and CD209b^25^. When VSIG4^+^ LPMs were depleted, VSIG4 expression was not regained for at least three weeks, and even at d56, the level of VSIG4^+^ LPMs did not yet reach steady-state levels, again suggesting a slow acquisition of the VSIG4 receptor. Finally, also the fact that VSIG4^+^ LPMs are somewhat less abundant in the PC of male mice could be in line with a slow VSIG4 acquisition, since LPMs in male mice have a higher turnover rate than in female mice^26^. Another factor that could influence VSIG4 expression is the presence of microbiota, whose metabolic products were shown before to affect the functionality of tissue-resident macrophages^13^. Interestingly, although the formation of VSIG4^+^ LPMs is not affected in germ-free conditions, their VSIG4 expression level is higher, suggesting that a tonic exposure to microbiota-derived molecules, either metabolic or structural, downregulates VSIG4 expression or increases its internalization.

However, if such a tonic triggering of VSIG4 in LPMs would exist, it does not lead to intracellular signaling and an altered gene expression. Indeed, we were unable to find any DEGs or DEPs to discern VSIG4^+^ from VSIG4^-^ LPMs at steady-state, suggesting that the VSIG4 receptor does not delineate a unique LPM subset. Nevertheless, VSIG4^+^ LPMs were superior phagocytic cells, implying that these cells could play a protective role during infectious and neoplastic disease in the PC. As a matter of fact, the specific depletion of VSIG4^+^ LPMs via our in-house generated anti-VSIG4-ADCC construct resulted in a higher first peak of parasitemia upon intraperitoneal *T. b. brucei* infection, as well as a decreased survival upon MC38 CRC inoculation in the PC.

The protective effect of VSIG4^+^ LPMs against peritoneal CRC tumor outgrowth is in line with findings demonstrating the LPM-mediated phagocytosis of metastasizing cancer cells in a model of ovarian cancer (ID8), whereby LPMs also play an important role in the early phases of cancer cell dissemination^27^. In patients with chronic liver disease that develop ascites, VSIG4^+^ macrophages were also found to be highly phagocytic as compared to VSIG4^-^ monocyte-derived macrophages^28^. In contrast, other studies claimed that LPMs support cancer cell metastasis and negatively affect anti-tumor T-cell immunity in cancer patients with peritoneal metastases^29,30^. Of note, VSIG4^+^ macrophages are also found in primary CRC tumors, where this marker mostly associates with C1Q, CD206 and CD163-expressing macrophages that display a tumor-promoting phenotype and whose presence has been associated with a worse outcome in several cancer types, including CRC and peritoneal metastasis^31–33^. It will be interesting to assess whether our anti-VSIG4-ADCC construct, or another approach that specifically depletes VSIG4^+^ cells, will be of therapeutic use against primary CRC tumors. However, such an approach holds the risk of increasing death due to peritoneal metastasis, as shown in this manuscript.

Altogether, our work sheds new light on the characteristics of VSIG4^+^ LPMs, identifying these cells as contributors to anti-parasitic and anti-metastatic responses in the peritoneal cavity.

## MATERIALS AND METHODS

### Mice

VSIG4^-/-3^, VISG4^+/+^, Flt3Cre^34^ and Rosa26-YFP^35^ mice were bred in-house. C57BL/6 mice were purchased from Janvier, France. Animal experiments were approved by the ethical commission for animal welfare at the Vrije Universiteit Brussel and performed in accordance with the guidelines set by the Belgian Council for Laboratory Animal Science.

### Cell culture and tumor models

The MC38 cell line was kindly provided by Massimiliano Mazzone (VIB-KULeuven, Belgium). The MC38 cell line was plated and cultured in DMEM supplemented with 10% FCS, Pen/Strep and glutamine at 37°C and 5% CO_2_. 10^6^ MC38-Thy1.1 cells were injected intraperitoneally in 200µL HBSS to mimic peritoneal metastasis of colorectal cancer cells.

### Trypanosoma infection

Bloodstream trypanosome parasites were stored at −80 °C as blood aliquots containing 50% Alsever’s solution (Sigma–Aldrich) and 10% glycerol (final v/v). Clonal pleomorphic *T. brucei* AnTat1.1E parasites were gifted by N. Van Meirvenne (Institute for Tropical Medicine, Belgium) and clonal *T. congolense* parasites (Tc13) were gifted by Dr. Henry Tabel (University of Saskatchewan, Canada). 6- to 12 - week-old female mice were infected with 5000 trypanosomes via intraperitoneal injection in a volume of 200 μL PBS. Parasite numbers in the blood were counted using a hemocytometer.

### Ex vivo preparation of single cell suspensions, flow cytometry and sorting

Processing of organs and subsequent flow cytometry analysis or sorting of cells is described in the supplementary data, including a list of antibodies used (Table S1).

### Phagocytosis assay

pHrodo Red *E. coli* Bioparticles (Thermofisher, P35361) and pHrodo Red *S. aureus* Bioparticles (Thermofisher, A10010) were prepared according to the manufacturer’s manual and injected intraperitoneally at 500µg/mL in 500µL HBSS. As a negative control, HBSS or unlabeled Bioparticles were injected (Thermofischer, S2859).

DiO cell labeling (Thermofischer, V22886) was prepared according to the manufacturer’s manual. MC38-Thy1.1 cancer cells were incubated with DiO dye (2.5µM) for 20 minutes at 37°C. The cancer cells were washed and spun down for 6 minutes at room temperature and 450G. The cancer cells were cocultured in sterile capped polystyrene tubes with peritoneal exudate cells in DMEM medium (Gibco) and a 37°C-incubator for 1 hour at 5% CO_2_. After incubation, the cells were washed with FACS buffer (1% FCS, 2mM EDTA) and prepared for antibody staining and flow cytometry analysis, as described in the supplementary materials and methods.

### Sample processing for CITE-sequencing

Peritoneal lavages were collected in 10mL ice-cold HBSS supplemented with 30µM actinomycin D. Three peritoneal lavages were pooled for each condition (HBSS-injected naive versus 18 hours post-*T. brucei brucei* infection). The single cell suspensions were stained with APC-Cy7-labeled anti-CD45 (30-F11), viability dye 7-Actinomycin D, TruStain FcX PLUS (Biolegend) and oligo-conjugated CITE-seq antibodies (Table S2) in FACS buffer (HBSS, 1% FCS and 2mM EDTA) supplemented with 3µM actinomycin D. After washing with FACS buffer, approximately 50.000 live CD45+ cells were sorted into FACS buffer (3µM actinomycin D) using the BD FACSAria III (BD Biosciences). The sorted cells were subsequently spun down and resuspended in PBS (0.04% BSA, 3µM actinomycin D) at an estimated final concentration of 1000 cells/µL. GEMs and CITE-seq libraries were prepared using a Chromium Next GEM Single Cell 3’ Gel Bead and Library kit, v3.1 as previously described^12^. Processing of CITE-sequencing data is described in the supplementary materials.

### Retrieval of published scRNAseq datasets

RNA expression matrices of a scRNA-seq dataset of naive CD102^+^ peritoneal macrophages of male and female mice were obtained from the Gene Expression Omnibus (GEO) database (GSE149014) (Bain et al.)^26^. Raw gene expression counts and cell metadata of a scRNA-seq dataset from tumors of 7 colorectal cancer cancer patients were downloaded from https://lambrechtslab.sites.vib.be/en/data-access (Qian et al)^36^. Processing of scRNA-seq datasets is described in the supplementary materials.

### Anti-VSIG4 constructs

The anti-VSIG4 Nanobody (Nb119) was obtained as previously described^37–38^. The anti-VSIG4 Nb sequence was cloned into a vector followed by a GlySer linker (10GS) and the WT or LALAPG-mutatetd Fc region of mouse IgG2a, resulting in the anti-VSIG4-ADCC and anti-VSIG4 ADCC-disabled constructs. The methods for generating, expressing, and purifying the anti-VSIG4 constructs is provided in the supplementary materials. VSIG4^+^ LPMs were depleted via intraperitoneal injection of 200 µg of anti-VSIG4-ADCC in 200µL HBSS. Injection regimens are shown in Figure 6A and 6C.

### In silico and ex vivo analysis of human CRC tissue

Human samples were obtained in accordance with the ethical guidance provided by the Declaration of Helsinki. Patients undergoing surgery for peritoneal metastasis provided written informed consent. The experimental protocol and informed consent documents were approved by the institutional review board (IRB) of Ghent University Hospital (ref. BC-06978). Fresh tumor tissue was obtained from 9 cancer patients with colorectal cancer metastasis to the omentum and mesentary. Surgical resection occurred at Ghent University Hospital (GI Surgery). Samples were transported on ice to the research facility at the Vrije Universiteit Brussel and processed for flow cytometry analysis. Single cell suspensions were prepared as previously described^39^. Correlation analysis of VSIG4 expression, CD163 and CD206 in tumors of colon adenocarcinoma patients was performed using http://gepia2.cancer-pku.cn/#index by Tang et al^40^.

### Statistics

Data is demonstrated as the mean ± standard error of the mean (SEM). GraphPad Prism 9.5.1 was used to calculate statistical significance. For pairwise comparisons, unpaired two-tailed Student’s t-test was performed. For the comparison of one or multiple groups, either a one-way analysis of variance (ANOVA) was performed with Tukey multiple comparisons test or a two-way ANOVA with Sidak multiple comparisons test was performed. Survival curves were analyzed using log-rank (Mantel-Cox) test. In case of statistically significant differences, the p value is shown on the graphs as such: *p≤0.05, **p≤0.01, ***p≤0.001, ****p≤0.0001.

Supplementary information is available at Cellular & Molecular Immunology’s website.

## Contributions

E.L: conceptualization, performed experiments, analyzed and visualized data and wrote the manuscript. S.M.A.: performed experiments and revised the manuscript. M.K., J.V.C., R.M.B., N.A.J., M.Z., N.A., Y.E., E.H.: performed experiments. D.K.: analyzed scRNAseq/CITEseq data. B.S., E.H., C.V., G.R. and D.L: conceptualization. J.A.V.G.: conceptualization, obtained funding support, supervised the study, wrote and revised the manuscript.

## Data availability

The data that support the findings of this study are available upon request from the corresponding author. CITE-seq data will be deposited in a public repository.

## Acknowledgements

We thank E. Omasta, M-T. Detobel, N. Van Riebeek, E. Vaneetvelde, M. Schuurmans, N. Abou, C. Papadopoulos and C. Stanley for technical and administrative assistance. We would like to thank J. Haustraete and his team at the VIB Protein Core for their support with Nb-Fc construct production and purification. E.L, S.M.A., P.M.R.B., A.A.C., M.K., N.A.J., R.M.B., and M.Z. were supported by an FWO predoctoral fellowship (1S67421N, 1S78122N, 1154722N, 1S23316N, FWOSB119, 1S18523N, 1199323N). J.V.C. was supported by a FWO-SBO (S001623N). S.A. was supported by a FWO research project (G090223N). E.L., S.M.A, M.K. and P.M.R.B. were supported by doctoral starter or finishing grants from Kom op Tegen Kanker. J.A.V.G, G.R., D.L., C.V., E.H, B.S., Y.E. and D.K. were supported by grants from FWO, Kom op Tegen Kanker, Stichting tegen Kanker and Vrije Universiteit Brussel.

Supplementary information is available at Cellular & Molecular Immunology’s website.

